# Optimized Protein–Excipient Interactions in the Martini 3 Force Field

**DOI:** 10.1101/2024.11.29.626008

**Authors:** Tobias M. Prass, Kresten Lindorff-Larsen, Patrick Garidel, Michaela Blech, Lars V. Schäfer

## Abstract

The high doses of drugs required for biotherapeutics, such as monoclonal antibodies (mAbs), and the small volumes that can be administered to patients by subcutaneous injections pose challenges due to high concentration formulations. The addition of excipients, such as arginine and glutamate, to high concentration protein formulations can increase solubility and reduce the tendency of protein particle formation. Molecular dynamics (MD) simulations can provide microscopic insights into the mode of action of excipients in mAb formulations but require large system sizes and long time scales that are currently beyond reach at the fully atomistic level. Computationally efficient coarse-grained models such as the Martini 3 force field can tackle this challenge but require careful parametrization, testing, and validation. This study extends the popular Martini 3 force field towards realistic protein–excipient interactions of arginine and glutamate excipients, using the Fab domains of the therapeutic mAbs trastuzumab and omalizumab as model systems. A novel all-atom to coarse-grained mapping of the amino acid excipients is introduced, which explicitly captures the zwitterionic character of the backbone. The Fab–excipient interactions of arginine and glutamate are characterized concerning molecular contacts with the Fabs at the single-residue level. The Martini 3 simulations are compared with results from all-atom simulations as a reference. Our findings reveal an overestimation of Fab–excipient contacts with the default interaction parameters of Martini 3, suggesting a too strong attraction between protein residues and excipients. Therefore, we reparametrized the protein–excipient interaction parameters in Martini 3 against all-atom simulations. The excipient interactions obtained with the new Martini 3 mapping and Lennard-Jones (LJ) interaction parameters, coined Martini 3-exc, agree closely with the all-atom reference data. This work presents an improved parameter set for mAb-arginine and mAb-glutamate interactions in the Martini 3 coarse-grained force field, a key step towards large-scale coarse-grained MD simulations of high-concentration mAb formulations and the stabilizing effects of excipients.

## Introduction

Monoclonal antibodies (mAbs) are successfully used across diverse targets and disease areas, such as asthma, infectious diseases, immunology, and oncology.^1^ Subcutaneous injection is the preferred route of administration for many antibody therapeutics for reasons that include self-administration for home medications, ease of use, reduction of hospitalization, and patient convenience. However, the high dosages required (approximately 1 to 10 mg of mAb per kg of body weight) and the small volumes that can be injected subcutaneously or into the muscle imply high-concentration formulations, typically greater than 100 mg/mL.^2^ Due to close intermolecular distances between mAbs, intermolecular interactions become non-negligible. Challenges arise due to non-ideal solution behavior at such concentrations, such as high viscosity, high opalescence, phase separation, aggregation or gelation, or the increased propensity for protein particle formation, which can negatively affect several steps during manufacturing and likely impair shelf-life of the drug product.^2–6^ Furthermore, high viscosity adversely affects the delivery of antibody therapeutics. This is because the injection needles of syringes are typically thin, ranging from 27 to 29 G (outer diameters of 0.42 mm to 0.32 mm, respectively) for subcutaneous injections. ^7^ This leads to increased tissue back pressure and high injection forces, which can compromise patient comfort. ^7–12^ In addition, aggregation poses risks to both efficacy and patient safety, including the potential for allergic responses and anaphylaxis.^13^

Current strategies for stabilizing protein drugs rely primarily on time- and resource-intensive trial and error methods. Such approaches often involve empirically derived heuristics to screen various combinations of buffers and excipients, including salts, amino acids, polyols, antioxidants, and surfactants. ^14,15^ One of the effective and frequently used amino acid-based excipients to mitigate protein aggregation and reduce viscosity are arginine salts.^16–18^ The mechanisms underlying the effect of arginine on solubility and aggregation are challenging to investigate, especially at the microscopic level. Molecular Dynamics (MD) simulations provide, in principle, the necessary resolution to obtain the detailed molecular insights desired to complement and interpret experimental results.^19,20^ Present MD studies investigating the effect of arginine on mAbs typically focus on the Fab domain. ^21–23^ Tilegenova et al. found in their MD simulations that aromatic protein residues are main interaction sites with arginine. ^21^ Golovanov et al. reported suppression of aggregation and improved protein solubility with glutamate as counterions to arginine. ^24^ In our previous atomistic MD simulation study on excipient interactions of the Fab domains of the therapeutic antibodies trastuzumab (Herceptin^®^) and omalizumab (Xolair^®^), we found preferential interactions of arginine with specific residues on the Fab surfaces (in particular charged and aromatic amino acids), and arginine-facilitated recruitment of glutamate towards the Fab surfaces.^23^ Omalizumab had predominantly stronger excipient interactions than trastuzumab. ^23^ These computational findings are in line with experimental long-term storage results ^23^ and shed light on the intricate intermolecular interactions that govern the macroscopic properties of protein–excipient formulations. Focusing on the Fab domains when studying the impact of excipients on mAbs is partly attributed to the significant influence of the Fab domains on protein–protein interactions, protein particle formation, viscosity, and solubility. ^25^ All-atom simulations of full-length mAbs in explicit solvent are computationally very expensive and therefore usually limited to single mAbs (monomers)^26–30^ or mAb dimers.^31^

Previously, we tackled this significant challenge and used multi-microsecond all-atom MD simulations of 4 mAb molecules in a simulation box to predict the viscosity of such high-concentration systems. ^32^ However, ideally simulation systems comprising of dozens of mAbs and for up to millisecond simulation times would be needed to generate the sampling required for statistically precise viscosity calculations, and also to enable the computational screening of a large number of systems and different formulation conditions. These requirements are currently out of reach for all-atom simulations but can be achieved with computationally more efficient coarse-grained (CG) models, which simplify molecular representations by combining groups of atoms into single beads. ^19,33^ These approaches range from high-level, where a single CG bead is a complete biopolymer, via one CG bead per amino acid residue models to low-level (i.e., detailed) models such as Martini.^34,35^ The widely used Martini model assigns fragments of 2 - 4 non-hydrogen atoms (including all covalently bonded hydrogens) to a single CG bead. Combined with its computational efficiency, the level of detail and chemical specificity retained in the CG-Martini model is beneficial for studying protein–excipient interactions.^36,37^ Especially with the release of Martini 3, which has improved protein–protein interaction parameters compared to its predecessor Martini 2,^38^ large-scale multi-protein systems became more accessible.^34,39,40^ At the same time, there is still room for improvement. On the one hand, Thomasen et al. observed too compact intrinsically disordered proteins (IDPs) and multidomain protein structures in Martini 3, which could be improved either by increasing the attractive protein-water interactions (by 10 %) or by decreasing the attractive protein–protein interactions (by 12 %).^41,42^ On the other hand, Pedersen et al. and Lamprakis et al. reported that protein–protein complexes that were found to be stable in all-atom simulations were unstable in Martini 3 simulations, suggesting underestimation of specific protein–protein interactions.^43,44^ These findings suggest that the underlying factors for the remaining imperfection of the model are more complex than simple global under- or overestimation of protein–protein or protein-water interactions. Concerning the explicit description of excipients, in the Martini force field amino acid-based excipients, such as arginine and glutamate, are modeled with the same CG beads (and corresponding parameters) that are used for the respective amino acid as part of a polypeptide chain, which we speculate could be a possible source of deviations.

For realistic large-scale CG-MD simulations of high concentration protein formulations (HCPFs), including the effect of excipients, an accurate description of all molecular interactions involved in such multicomponent systems is needed, i.e., protein–protein, protein– excipient, excipient–excipient, etc, and also the interactions with the solvent (water) and mobile ions. The delicate balance between all these contributions governs the properties of the solution. Previous Martini force field development work has primarily focused on protein–protein and protein–water interactions (as described above), but less on improving the description of excipients. Thus, a key component required for MD simulations of HCPFs with the Martini force field is missing. In this work, we investigated the protein– excipient interactions in Martini 3 of arginine (Arg) and glutamate (Glu) excipients with the Fab domains of the therapeutic mAbs trastuzumab and omalizumab. First, we propose a novel all-atom to CG mapping in which two charged CG beads are assigned to the amino acid backbone of the excipient molecule (following the building block principle of Martini 3), thereby explicitly capturing the zwitterionic nature of the individual (free) amino acid excipient molecules. Then, we compute the Fab–excipient interactions of arginine and glutamate for different excipient systems, 150 mM Arg/Cl and 150 mM Na/Glu (protein-to-excipient molar ratio of 1:90). The molecular contacts of the excipients to the trastuzumab Fab domain surface residues are compared with results from all-atom simulations with the CHARMM36m-NBF force field that includes the recent refinement of cation–*π* interactions of Liu et al. ^45^ The molecular contacts of the excipient molecules with the protein amino acid residues (or residue types) in Martini 3 vs. CHARMM36m-NBF are used to optimize the protein–excipient interaction parameters in the Martini 3 force field. Finally, the improved parameter set is verified with simulations of a different mAb that was not used in the parametrization (training) procedure, omalizumab, and with additional simulations of trastuzumab in an equimolar excipient mixture of 75 mM Arg/Cl and 75 mM Na/Glu. Overall, this work presents a straightforward approach to optimize protein–excipient interactions in the Martini 3 coarse-grained force field and an improved Martini 3 parameter set for describing mAb–arginine and mAb–glutamate interactions.

## Methods

### Setup of Simulation Systems

#### Atomistic systems

The atomic coordinates of the omalizumab Fab domain were obtained from the protein data bank (PDB: 2XA8).^46^ For trastuzumab, the structure of the D185A light chain mutant (PDB: 6BHZ)^47^ was used, in which missing residues were added and the alanine in the light chain at position 185 was replaced with aspartate using SWISS-MODEL and Swiss-Pdbviewer.^48^ The titratable amino acid side chains were modeled in their default protonation states corresponding to pH 7 conditions, as assigned by the GROMACS tool pdb2gmx. His207 and His208 in the heavy chains of omalizumab and trastuzumab, respectively, were modeled in their charged (doubly protonated) states, while all other histidine residues were singly protonated. Each Fab domain was placed in a cubic simulation box with ca. 10 nm edge length. Arginine and/or glutamate molecules were introduced at the desired concentrations. To neutralize the net charge of the simulation boxes due to the introduction of the positively charged arginine and negatively charged glutamate excipients, corresponding amounts of chloride and sodium ions were added. Furthermore, trastuzumab Fab was also simulated in an equimolar mixture of 75 mM arginine and 75 mM glutamate (including 75 mM sodium and 75 mM chloride ions to maintain consistent ionic strength across all three systems). The remaining net charge in each simulation box was neutralized by adapting the number of chloride and sodium ions.

#### Coarse-grained systems

The coarse-grained models of the Fab domains of omalizumab and trastuzumab were generated with the Martinize 2 script ^49^ using the atomistic simulation systems described above as input and applying side chain corrections proposed by Herzog et al. ^50^ For the all-atom to coarse-grained conversion, we defined updated mapping instructions for the arginine and glutamate excipients, representing the backbone with two beads instead of one (which is the default in Martini), see Figure 1A. Not only does this approach capture the zwitterionic character of the backbone, but the molecular surfaces also match those of all-atom models much closer than the CG-Martini models of the amino acid residues with single backbone beads (Figures 1B and 1C). The C_*α*_ and ammonium group were assigned to a Martini SQ4p bead type, while the carboxylate group was assigned to an SQ5n bead (Figure 1A), following the recommended Martini 3 mappings for these functional groups. ^39^

**Figure 1:**
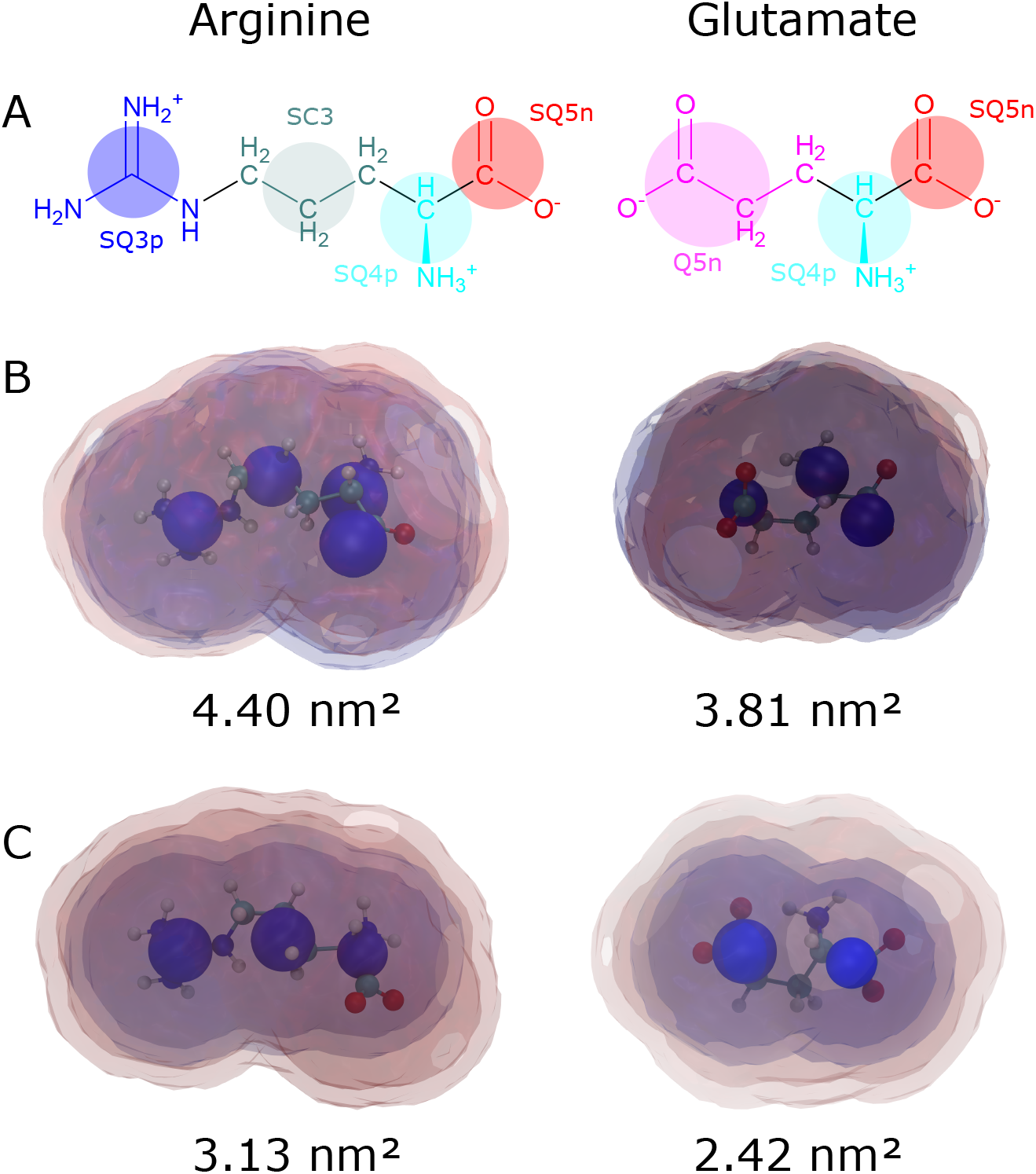
Coarse-grained arginine (left column) and glutamate (right column) models in comparison with the all-atom structures. A) Lewis structures of the excipients and schematics of the mapping of the atoms to the coarse-grained (CG) beads. B) and C) Connolly surfaces (transparent blue surfaces) and Martini 3 CG beads (blue spheres). The red volumetric objects show the Connolly surfaces of the underlying atomistic structures. The coarse-grained models presented in B) are the new CG models with two backbone beads developed in this work, while the models shown in C) are the default Martini 3 representations with a single backbone bead. The numbers below the 3D renderings denote the solvent-accessible surface area (SASA) of the CG structures. The SASAs of the corresponding atomistic arginine and glutamate structures are 4.68 nm^2^ and 3.86 nm^2^, respectively.

### Molecular Dynamics Simulations

The MD simulations in this work were carried out with GROMACS (version 2021.1). ^51^ For the all-atom simulations, the CHARMM36m-NBF force field with improved cation–*π* interactions^45,52,53^ was used. For the present study, cation–*π* interactions are important for contacts of arginine excipients with aromatic amino acid side chains. For water, the CHARMM version of the TIP3P water model was used. Short-range electrostatic and Lennard-Jones interactions were computed up to a distance cutoff of 1.2 nm between atom pairs. Forces were smoothly switched to zero between 1.0 and 1.2 nm. Long-range electrostatic interactions were computed with the smooth particle-mesh Ewald (PME) method^54^ with default settings in GROMACS. After energy minimization, the systems were equilibrated for 2 ns with position restraints on the proteins at a constant temperature of 298 K and constant pressure of 1 bar, implemented with the velocity rescaling thermostat with a stochastic term^55^ and the stochastic cell rescaling barostat,^56^ respectively (with 0.1 and 2.0 ps coupling time constants, respectively). The LINCS^57^ algorithm was used to constrain the lengths of covalent bonds within the protein involving hydrogen atoms, and SETTLE^58^ was used to constrain the water molecules. The equations of motion were integrated with 2 fs time steps. Finally, for each simulation system, five independent production MD simulations were performed for 500 ns each, initiated with different random seeds for generating the initial atomic velocities. These simulation trajectories were taken from our previous work. ^23^

The corresponding coarse-grained simulations employed the Martini 3 force field ^39^ with the new excipient models introduced in this work. The same protocol as mentioned before was followed, except for equilibrating for 20 ns and production runs of 1 *µ*s. The recommended “new-rf”^59^ simulation settings were used, which included the use of a 1.1 nm nonbonded cutoff with reaction field electrostatics and 20 fs integration time steps. The Fab structures were restrained with an elastic network between backbone beads using the default settings of Martinize 2 (elastic bond force constants of 500 kJ mol^−1^ nm^−2^ and lower and upper cutoffs of 0 nm and 0.9 nm, respectively). Temperature and pressure were kept constant at 300 K and 1 bar, respectively (with the same thermostat and barostat as in the all-atom simulations). For statistical analysis, the first 100 ns were discarded and the remaining 900 ns were split into three blocks of 300 ns.

### Parametrization of Martini 3 Arginine and Glutamate Excipient Models

The intramolecular interaction parameters were generated using all-atom simulations (mapped to CG coordinates) as a reference. To that end, three cubic simulation boxes with 8 nm edge length were generated and filled with 1) 150 mM Arg/Cl, 2) 150 mM Na/Glu, and 3) 75 mM Arg/Cl + 75 mM Na/Glu, each solvated with water. MD simulations were performed for 100 ns using the CHARMM36m-NBF force field^45^ (at constant temperature of 298 K and pressure of 1 bar). The trajectory was saved to disk with 10 ps time spacing and mapped to the CG representation with the Martinize 2 script. ^49^ The bond and angle distributions were used to initially estimate and then adjust the bond and angle parameters of the CG excipient models. Then, Martini 3 simulations analogous to the excipient-only all-atom simulations were performed to first, validate the final bond and angle distributions against the corresponding ones from the all-atom reference simulations, and second to compute the excipient–excipient radial distribution functions (RDFs). The intermolecular Lennard-Jones (LJ) interactions between excipients were scaled in Martini 3 to match the integrals of the RDFs (as a measure of the effective interaction strength) obtained with the CHARMM36m-NBF force field.

### Parametrization of Protein–Excipient Interactions in Martini 3

A key analysis performed in this work is based on molecular contacts between excipient molecules and protein surface residues. For each frame of the MD trajectories, whenever at least one atom (or bead, for the CG simulations) of a given excipient molecule was within a cutoff distance of at least one atom (or bead) of a particular protein surface residue, one molecular contact was counted. These contacts were then averaged over all trajectory frames. The cutoff distances used for contact counting were 0.50 nm and 0.75 nm in the atomistic and Martini 3 simulations, respectively. The different cutoff values are required due to the different resolutions of the two models. To verify the direct comparison of contacts obtained with 0.50 nm and 0.75 nm cutoffs used in the analyses of the all-atom and coarse-grained simulations, respectively, the all-atom trajectory was mapped to the Martini 3 representation and the contacts were compared (Figure S1). The Pearson correlation coefficients between the contacts directly computed from the all-atom trajectory (with 0.50 nm cutoff) and the ones from the same trajectory analyzed in terms of its CG coordinates (with 0.75 nm cutoff) are *r* = 0.98 for arginine and *r* = 0.96 for glutamate (with slopes of the linear fits of 0.995 and 0.989, respectively), showing that the two cutoff values provide highly similar results.

To adjust the protein–excipient interactions in the Martini 3 force field, we scaled the well depths *ϵ*_res-exc_ of the LJ 12-6 interaction potentials between the protein residues and the excipients. In Martini 3, the pairwise LJ interactions between all bead types are explicitly defined in an interaction matrix. Since the residue–excipient interactions are to be reparametrized for each residue type separately, we expanded the Martini 3 interaction matrix by adding residue type- and excipient type-specific bead types, whose LJ parameters were copied from their respective default values. The new interaction parameters between the amino acid residues and the excipients were then calculated via

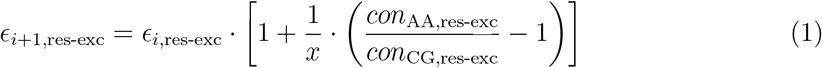

Here, *ϵ*_*i*,res-exc_ is the depth of the LJ potential well. The indices “*i*”, “res”, and “exc” denote the iteration counter, residue type and excipient, respectively. The number of contacts is indicated by “*con*”, where “AA” and “CG” correspond to all-atom and coarse-grained simulations, respectively. The “learning rate” *x* is introduced to avoid too strong scaling in subsequent iterations, with its value chosen between 10 and 20. A value of *x* = 10 was employed during the early iterations, with adjustments in subsequent iterations depending on the difference of contact counts in the Martini 3 and CHARMM36m-NBF simulations. In the above scaling procedure, we did not differentiate between the different beads within a given protein residue or excipient, that is, the same scaling of LJ interaction strength was applied to all beads of that residue type or excipient. Removing this restriction would enable a more flexible fitting procedure at the expense of increasing the dimensionality of the parameter space. In light of the results obtained with the simpler strategy adopted in this work (see below), we leave this possible extension for future work. We used the protein– excipient contacts from the simulations with trastuzumab as reparametrization target and the simulations with omalizumab for validation. The statistical errors of the molecular contacts were estimated using the standard deviations between the five 500 ns trajectories for the all-atom simulations, and the standard deviations between the three 300 ns trajectory blocks for the CG simulations.

## Results and Discussion

### Excipient Parametrization

Here we introduce new Martini 3 models of arginine and glutamate excipients with two CG beads that describe the backbone of these free amino acids. This approach involves the definition of a new AA-to-CG mapping scheme and the use of different bead types (Figure 1). It has two main advantages. First, the zwitterionic character of the backbone is explicitly represented by the positively and negatively charged beads for the N- and C-terminus, respectively. Second, as shown in Figure 1, the solvent-accessible surface areas (SASAs) of the excipients of 4.40 nm^2^ (arginine) and 3.81 nm^2^ (glutamate) match the all-atom SASAs (4.68 nm^2^ and 3.86 nm^2^, respectively) much closer than the models with a single backbone bead (3.13 nm^2^ and 2.42 nm^2^, respectively). The finding that the CG-Martini model with a single backbone bead does not represent the molecular surfaces well is not surprising in light of the fact that it was developed for describing amino acid backbones as part of polypeptide chains, and not for isolated amino acids with two charged termini. The bonded (intramolecular) force field parameters of the new CG models for arginine and glutamate were generated with atomistic simulations as reference (see Methods). The CG and all-atom distributions are plotted in Figures S2 and S3 and the final intramolecular parameters are listed in Table 1. For the majority of the cases, the bond and angle distributions from the atomistic excipient-only simulations closely agree with the corresponding CG distributions, thus the bonded parameters were only slightly adjusted.

**Table 1:** Intramolecular interaction parameters for the adapted Martini 3 models of arginine and glutamate. BBO and BBN are the SQ5n and SQ4p beads for the backbone carboxyl and amino-groups, respectively, and SC1 (and SC2, for arginine) refer to the side chain beads (Figure 1A). The parameters *r*_*eq*_, *θ*_*eq*_, *k*_*bond*_, and *k*_*θ*_ are the equilibrium bond length and angle and the force constants of the harmonic bond and angle potentials, respectively.

|  | Arginine |  | Glutamate |  |
| --- | --- | --- | --- | --- |
| Beads | Bonds |  |  |  |
| | $r_{eq}/nm$ | $k_{bond}/\frac{kJ}{mol \cdot nm^2}$ | $r_{eq}/nm$ | $k_{bond}/\frac{kJ}{mol \cdot nm^2}$ |
| BBO-BBN | 0.251 | 99103 | 0.252 | 99103 |
| BBN-SC1 | 0.293 | 7647 | 0.331 | 991 |
| BBO-SC1 | 0.377 | 1810 | 0.468 | 2143 |
| SC1-SC2 | 0.354 | 5119 | — | — |
|  | Angles |  |  |  |
| | $\theta_{eq}/deg$ | $k_{\theta}/\frac{kJ}{mol \cdot rad^2}$ | $\theta_{eq}/deg$ | $k_{\theta}/\frac{kJ}{mol \cdot rad^2}$ |
| BBO-BBN-SC1 | 50.722 | 146.3 | 42.210 | 53.690 |
| BBO-SC1-SC2 | 139.833 | 33.5 | — | — |
| BBN-SC1-SC2 | 132.656 | 15.5 | — | — |

After having optimized the intramolecular interaction parameters of the excipients, we performed CG-MD simulations with only excipients (no proteins). Figure 2 shows the excipient–excipient RDFs obtained from these simulations. The dashed orange lines show the RDFs with the original Martini 3 parameters (that is, before adjusting the intermolecular interaction strengths), which exhibit reduced attraction compared to the atomistic reference data obtained with the CHARMM36m-NBF force field. The backbones of the excipients consist of SQ4p and SQ5n beads in the CG model, which are among the most polar Martini 3 beads. We decided to specifically adjust the excipient–excipient and protein–excipient interactions. This parametrization was achieved by globally rescaling the Lennard-Jones well depths of all involved molecule-specific bead types, see equation 1. To match the coarse-grained and all-atom results for the excipients, the LJ interaction strengths in the Martini 3 force field between the arginine excipients (arginine–arginine) were increased by 65 %, and those between glutamate excipients (glutamate-glutamate) were increased by 40 %. The area under the curve of the Arg–Arg RDF obtained with this reparametrized model (Figure 2, orange line) agrees with the area under the CHARMM36m RDF, which is plotted in blue in Figure 2.

**Figure 2:**
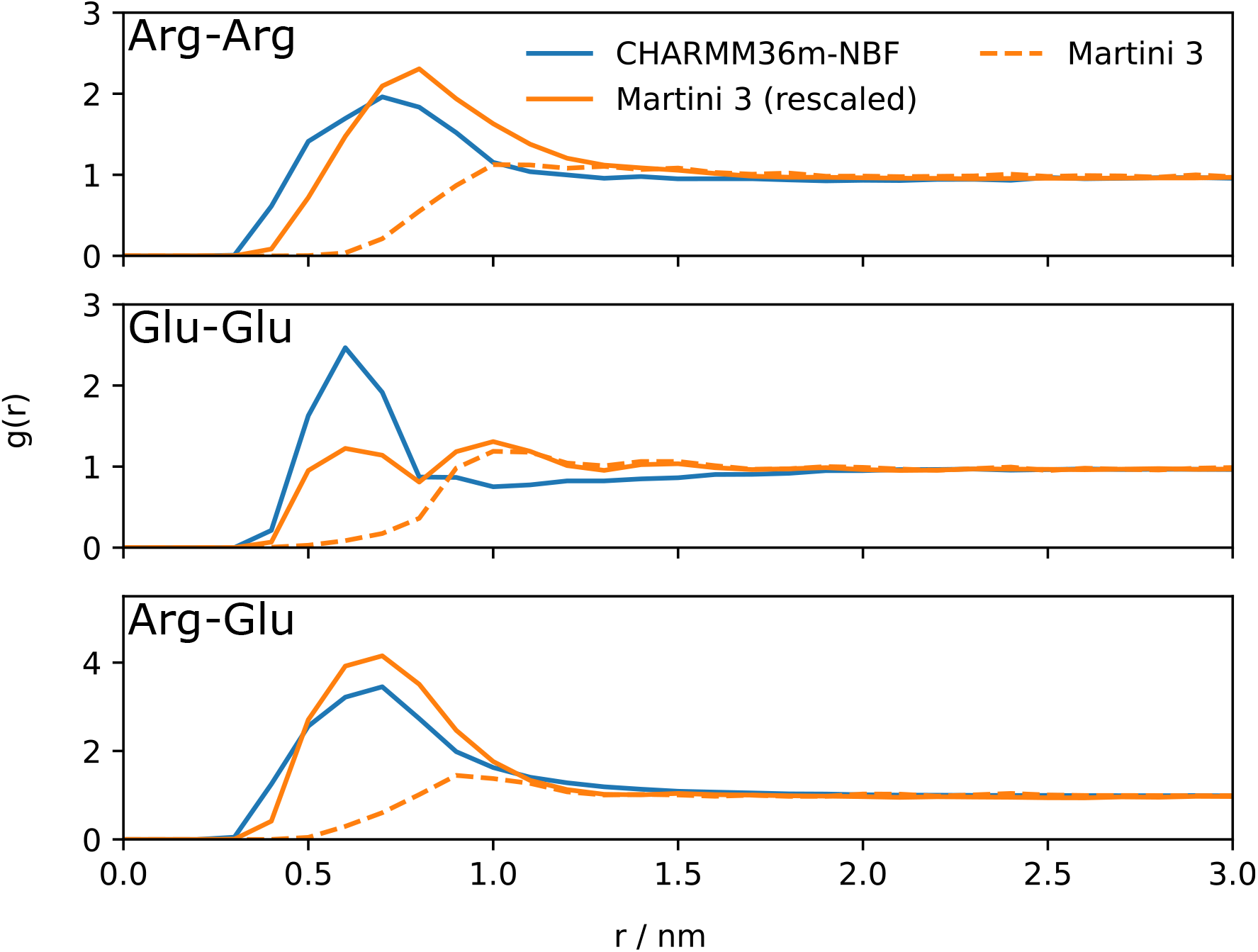
Radial distribution functions of the centers of mass of the excipients. The graphs from top to bottom were obtained from the 150 mM Arg/Cl, 150 mM Na/Glu and 75 mM Arg/Cl + 75 mM Na/Glu simulations, respectively.

Notably, in both the original and the reparametrized Martini 3 force fields, the Glu– Glu RDF (Figure 2, middle panel) has two maxima, corresponding to a direct contact pair and a solvent-separated configuration at ca. 0.6 nm and 1.0 nm, respectively. Adjusting the LJ well-depths leads to the maximum at 0.6 nm, which has lower intensity than the CHARMM36m-NBF maximum at this distance. However, after reparametrization, the areas below the atomistic and coarse-grained RDFs are similar, so we decided not to increase the Glu–Glu interactions further. We speculate that the additional maximum at 1.0 nm might be due to packing effects related to the 3-bead model of glutamate. Notably, after the reparametrization of the Arg–Arg and Glu–Glu interactions described above, Arg-Glu rescaling was not necessary to obtain agreement with the atomistic reference data (bottom panel of Figure 2).

### Martini 3 Before Protein–Excipient Reparametrization

After introducing the new zwitterionic backbone CG-Martini 3 models of arginine and glutamate excipients and parametrizing the excipient–excipient interactions in Martini 3, we simulated the trastuzumab Fab domain with two different excipient conditions, 150 mM Arg/Cl and 150 mM Na/Glu. Preferential excipient–residue interactions were analyzed by counting molecular contacts. In the atomistic simulations, a molecular contact was counted if at least one excipient atom was within 0.50 nm of one or more atoms of a protein residue. In the CG simulations, this cutoff value for the bead-bead contacts was increased to 0.75 nm (Figure S1). The results obtained with the new Martini 3 excipient models (but the original unscaled LJ interactions as defined in Martini 3) are compared to the all-atom simulations with the CHARMM36m-NBF force field in Figure 3.

**Figure 3:**
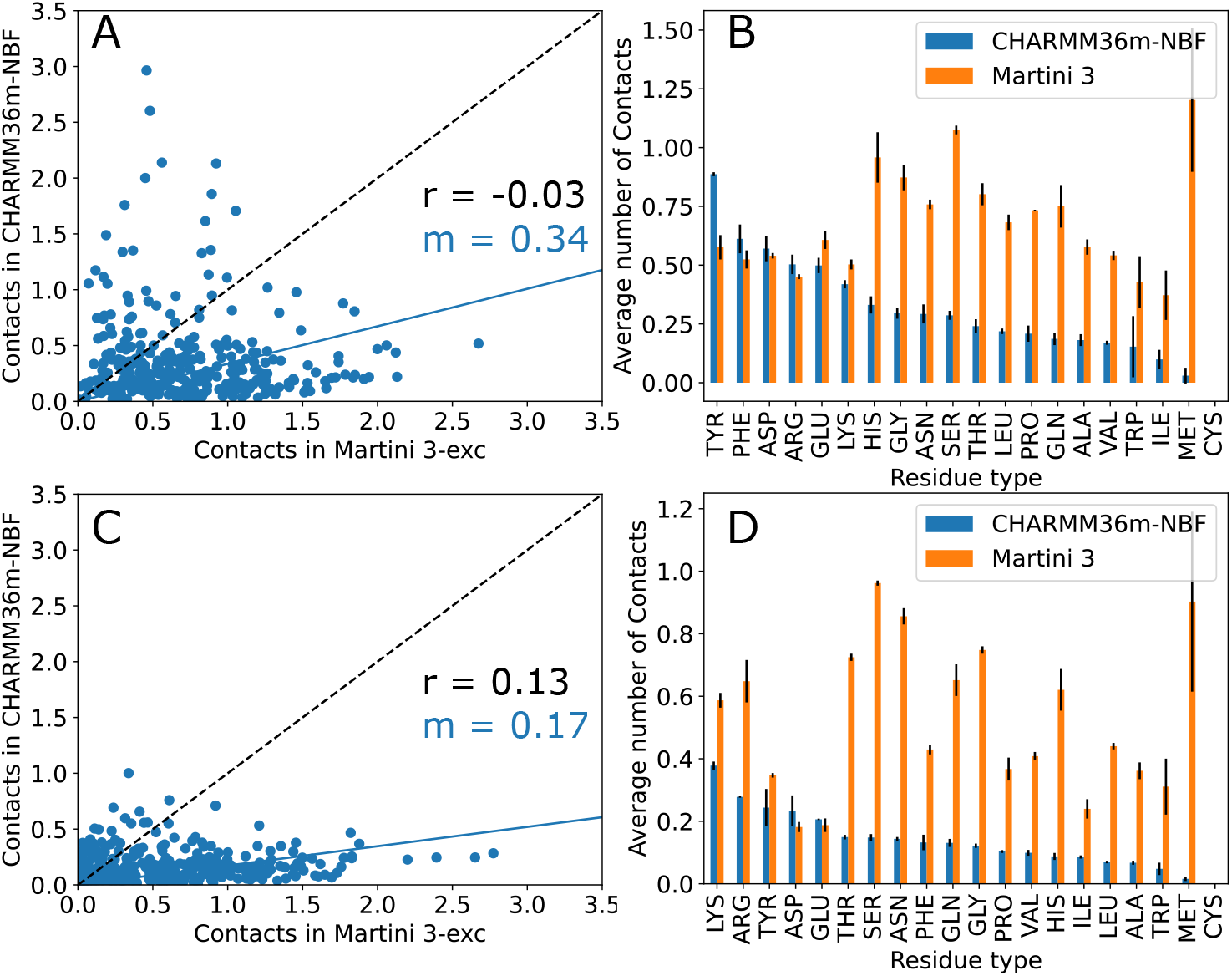
Comparison of residue–excipient contacts for the trastuzumab Fab domain with CHARMM36m-NBF and unscaled Martini 3. The residue–arginine contacts are shown in A and B, the residue–glutamate contacts in C and D. In the correlation plots (A and C), the residue–excipient contacts with Martini 3 and CHARMM36m-NBF are compared (r denotes the Pearson correlation coefficients, m the slope of the linear fits (blue lines)). The dashed black lines are the diagonals (*y* = *x*). The average number of excipient–residue contacts with CHARMM36m-NBF and Martini 3 are shown in B and D for the different amino acid residue types. The error bars in panels B and D represent the standard deviations.

Figure 3A shows the correlation plot of trastuzumab Fab–arginine molecular contacts in both force fields. With a Pearson correlation coefficient (PCC) of *r* = −0.03, Martini 3 and CHARMM36m-NBF do not correlate. On average, the number of molecular contacts observed in Martini 3 is about two to three times higher than in the all-atom simulations, suggesting a systematically too strong attraction between protein surface residues and excipients. Furthermore, the specificity seen in the different high-contact regions differs significantly, especially concerning the distinct pattern of preferential excipient interactions seen in the complementarity-determining regions^23^ (see Figure S4 for a detailed representation of the protein–excipient interactions along the Fab sequence). This is also reflected in the overestimation of the residue type-averaged contacts with arginine (Figure 3B). The general trend of preferable interactions of the trastuzumab Fab domain with arginine excipients significantly differs between the all-atom and the coarse-grained force fields. While the average local interactions of arginine excipients with Phe, Asp, Arg, Glu and Lys amino acid residues match between Martini 3 and CHARMM36m-NBF, they are underestimated for Tyr residues and overestimated for other amino acid residue types. Furthermore, the all-atom simulations revealed on average significantly fewer contacts of glutamate excipients with the trastuzumab Fab domain compared to arginine (compare upper and lower panels in Figure 3). The glutamate contacts in Martini 3 compared to CHARMM36m-NBF are presented in Figures 3C and 3D, showing a similar overestimation of the molecular contacts in the original Martini 3 interaction parameters as seen for arginine excipients. Furthermore, with the original interaction strengths, the number of Fab-glutamate contacts is as high as the Fab-arginine contacts, in strong contrast to the all-atom simulations in which a preferential interaction of the Fab domain with arginine excipients (over glutamate) is observed. Therefore, for a more accurate description of specific protein–excipient interactions, we concluded that it is necessary to adjust the protein–excipient interaction parameters in Martini 3.

### Reparametrized Martini 3-exc Force Field

The differences in excipient–residue contacts shown in Figures 3B and D serve as a target to adjust the Martini 3 interaction parameters. In Martini 3, the beads used to describe amino acid residues in proteins are assigned to represent the key chemical fragments (or “building blocks”) of each residue. Some of these fragments are represented by the same bead type in Martini 3, for example, Phe, Tyr, and Trp have TC5 type beads describing the aromatic ring. To reparametrize the excipient–residue interactions we defined residue-specific and excipient-specific bead types with, at the beginning of the parametrization procedure, the same LJ interaction parameters as in the original Martini 3 force field. Then, the LJ parameters were adjusted iteratively to match the molecular contacts of the CHARMM36m-NBF data (see Methods). The final results (after 15 iterations, equation 1) obtained with the reparametrized Martini 3 force field, coined Martini 3-exc, are presented in Figure 4 and the final scaling factors of the LJ well-depths are listed in Table 2.

**Table 2:** Scaling factors of the Lennard-Jones well-depths between protein residues and excipients in Martini 3-exc. Residue types that occur fewer than 5 times at the surface of trastuzumab are marked with an asterisk.

| Residue | Arginine Scaling Factor | Glutamate Scaling Factor |
| --- | --- | --- |
| Ala | 0.85243063 | 0.78866005 |
| Arg | 1.42525544 | 1.05145992 |
| Asn | 0.88332562 | 0.78235118 |
| Asp | 1.17903206 | 1.43715351 |
| Gln | 0.92320811 | 0.90999592 |
| Glu | 1.14044223 | 1.43277269 |
| Gly | 0.82831058 | 0.80240124 |
| His* | 0.65421264 | 0.52236041 |
| Ile* | 0.67424864 | 1.06553751 |
| Leu | 0.91315563 | 0.88385436 |
| Lys | 1.14728312 | 1.05905944 |
| Met* | 0.50578418 | 0.46673367 |
| Phe | 1.32787235 | 1.24383757 |
| Pro | 0.82090591 | 1.03344047 |
| Ser | 0.80059120 | 0.73218966 |
| Thr | 0.82067907 | 0.89391894 |
| Trp* | 0.49368134 | 0.59695372 |
| Tyr | 1.17262296 | 1.14480006 |
| Val | 0.88504739 | 0.97775697 |

**Figure 4:**
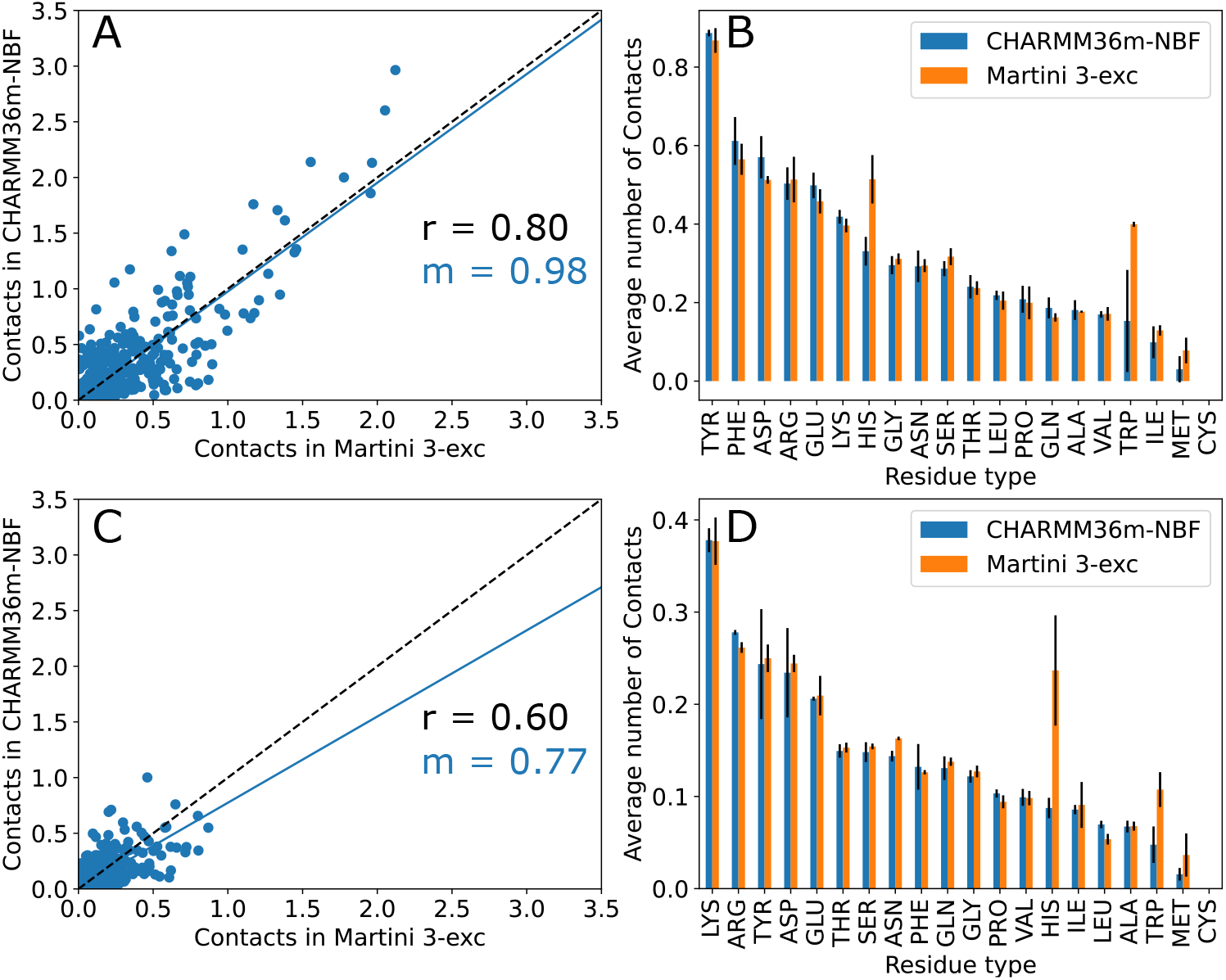
Residue–excipient contacts from the Martini 3-exc simulations of the trastuzumab Fab domain with reparametrized residue–excipient LJ interaction strengths are compared to the CHARMM36m-NBF results. The residue-arginine contacts are shown in A and B, and the residue-glutamate contacts in C and D. In the correlation plots (A and C), the residue– excipient contacts with Martini 3-exc are compared to CHARMM36m-NBF (r denotes the Pearson correlation coefficient, m is the slope of the linear fits (blue lines)). The dashed black lines are the diagonals (*y* = *x*). The average number of excipient–residue type contacts are shown in B and D with error bars presenting the standard deviations.

Figure 4A depicts the arginine contact correlation plot of CHARMM36m-NBF with the Martini 3-exc simulations. The PCC of the arginine contacts between both models is *r* = 0.80, which is significantly higher compared to the original Martini 3 force field (Figure 3). The arginine–residue type contact comparison in Figure 4B also highlights the major improvements achieved with the new interaction parameters. Only the arginine contacts with His and Trp residues still deviate from the all-atom reference values. However, these deviations have to be interpreted with care due to the low number of His and Trp residues at the surface of trastuzumab (4 and 3 residues, respectively), which implies limited statistics.

Similar to arginine, also for glutamate the agreement between the atomistic and coarse-grained simulations is considerably improved (Figures 4C and 4D), with a PCC of 0.60 compared to 0.13 with the original Martini 3 interactions strengths. The smaller correlation found for the glutamate–residue contacts can be explained by the smaller range of the values. The average number of contacts of trastuzumab residues with arginine is 0.38, with a broader range of values and a significant fraction of data points above 1.5 (Figure 4A), whereas the average number of glutamate contacts is only 0.13. Therefore, small differences between the all-atom and coarse-grained data have a larger effect on the correlation of the glutamate contacts. In Figure 4D the glutamate–residue type contacts are compared between the Martini 3-exc and CHARMM36m-NBF simulations. As discussed above for arginine, the His and Trp contacts in Martini 3-exc are still overestimated, which did not improve with more iterations (equation 1). This is likely related to the low number of His and Trp surface residues in the trastuzumab Fab domain (4 and 3, respectively).

The majority of arginine–residue type contacts in our reparametrized Martini force field reached good agreement with the atomistic data after scaling down the arginine–residue type interactions (Table 2), that is, making them less attractive (reduced well-depth of the LJ potential). In contrast, arginine interactions with Arg, Asp, Glu, Lys, Phe, and Tyr were increased to match the CHARMM36m-NBF data. The same trend is observed for the interactions of glutamate excipients. At first glance, the necessity to increase selected excipient–residue interactions seems counterintuitive, because the contacts shown in Figure 3 are overestimated for all residue types (except Tyr and Phe). However, excipient–residue interactions cannot be attributed to individual residues alone, but they are also affected by corecruitment due to excipient interactions with neighboring amino acid residues on a local patch on the protein surface.^23^

The residue types in Table 2 marked with asterisks are rare on the trastuzumab Fab surface (less than 5 surface residues of the respective type, none in case of Cys). Therefore, the limited sampling in terms of the local surface patch context of these residues might influence the average number of excipient–residue contacts, implying that the resulting interaction parameters are unlikely to be applicable to significantly different proteins. In particular, the interaction strengths of the excipients with His and Trp were scaled down, but still overestimate the molecular contacts compared to CHARMM36m-NBF (Figures 4B and 4D).

The results presented here were obtained by optimizing the excipient–Fab interactions for trastuzumab. In total, 16 iterations with varying *x* in Equation 1 were performed, which systematically improved the agreement between the coarse-grained and all-atom force fields step by step. The improvements between consecutive refinement iterations decrease steadily, which means that already after a few (e.g., 4 or 5) iterations markedly improved results can be obtained. Overall, the new Martini 3-exc force field parameters yield molecular contacts for trastuzumab that closely match the all-atom reference data. The LJ interaction strength reparametrization presented in this work was carried out for the trastuzumab Fab domain, and generalizations about protein–excipient interactions in Martini 3 should be made with caution. However, we expect that Martini 3-exc will also yield an improved description of the excipient interactions for other mAbs due to the similarity of antibodies, especially in the consensus constant regions of the same subclass. For more distant sequences, we expect that our model constitutes an improved starting point for additional fine tuning using the approach we have presented.

### Equimolar Excipient Mixtures

The arginine–excipient and glutamate–excipient parameters of Martini 3-exc were optimized with single excipient systems with chloride (for arginine) or sodium (for glutamate) counterions, respectively. To test for transferability, simulations with 75 mM concentrated equimolar mixtures of arginine and glutamate excipients (with 75 mM Na/Cl) were performed and the molecular excipient–residue contacts compared between Martini 3-exc and CHARMM36m-NBF.

Figure 5 presents the respective correlation plots. The molecular contacts of arginine, (Figure 5A) demonstrate a high level of consistency between the two force fields, with a strong correlation (*r* = 0.82). Consequently, the new arginine–residue interactions in Martini 3 are expected to be robust toward changes in both arginine excipient concentration and (minor) changes in the solvation environment. The molecular contacts of glutamate (Figure 5) have a PCC of *r* = 0.36 and are more strongly affected by the changes of the solvation conditions. Due to the low number of molecular contacts of glutamate to trastuzumab, this result supports the notion that excipients with comparatively weak protein interactions are more influenced by changes in the environment. However, when averaged over all trastuzumab residues, the Martini 3-exc glutamate interactions are reasonably close to those obtained from the all-atom reference simulations with average contact counts of 0.05 and 0.09, respectively.

**Figure 5:**
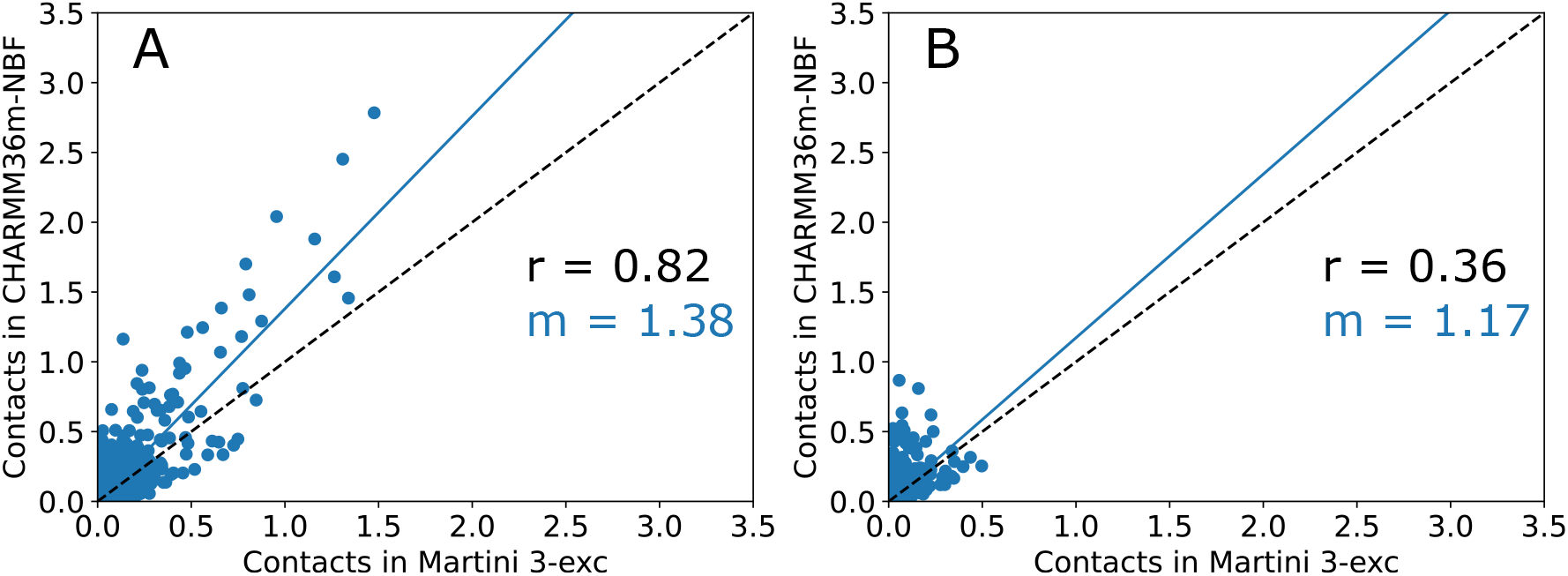
Correlation of protein–excipient contacts between CHARMM36m-NBF and Martini 3-exc simulations of the trastuzumab Fab domain with equimolar concentrations of 75 mM Arg/Cl and Na/Glu (r denotes the Pearson correlation coefficient, m is the slope of the linear fits (blue lines)). The dashed black lines are the diagonals (*y* = *x*). The arginine– residue and glutamate–residue contacts are depicted in panel A and B, respectively.

### Crossvalidation of Martini 3-exc for a Different Antibody

To test the transferability of the new Martini 3-exc parameter set to other antibody Fab domains that were not included in the parametrization procedure, we simulated the omalizumab Fab domain with the excipients arginine and glutamate. As described above for trastuzumab, we analyzed the molecular contacts by comparing the results to all-atom simulations carried out with the CHARMM36m-NBF force field. The omalizumab and trastuzumab Fab domains have sequence identities of 82 % and 91 % for the heavy and light chains, respectively. The correlation plots of excipient–residue contacts between CHARMM36m-NBF and Martini 3-exc are presented in Figure 6 for both arginine (Figure 6 A) and glutamate (Figure 6 B). The corresponding correlation plots with the unscaled Martini 3 force field are shown in Figure S5.

**Figure 6:**
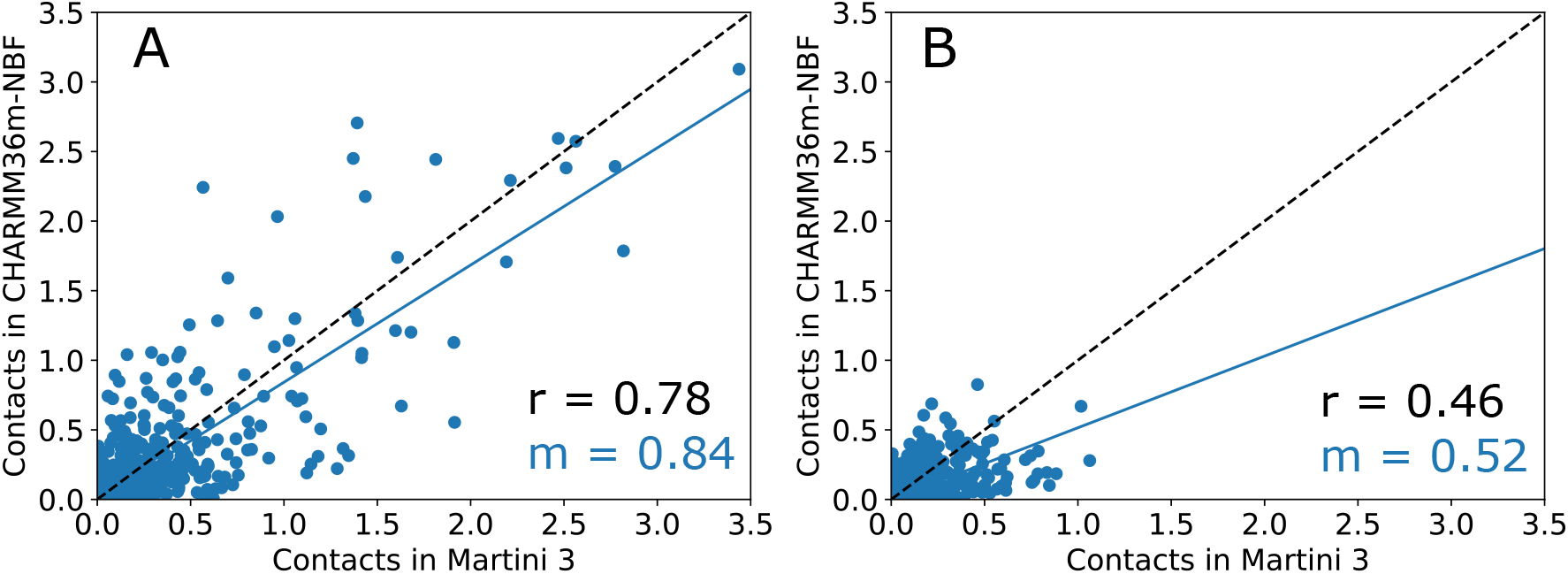
Correlation of protein–excipient contacts between CHARMM36m-NBF and Martini 3-exc simulations of the omalizumab Fab domain (r denotes the Pearson correlation coefficient, m is the slope of the linear fits (blue lines)). The dashed black lines are the diagonals (*y* = *x*). The arginine–residue and glutamate–residue contacts are depicted in panels A) and B), respectively.

There is a high correlation in the number of arginine–residue contacts between reparametrized Martini 3-exc and the CHARMM36m-NBF force field (Pearson correlation coefficient of 0.78). The slope of 0.84 also highlights a close match between overall interaction strengths between both contact data sets. We conclude that the parametrization strategy works well for arginine excipients and that the Martini 3-exc parameters can provide more accurate results also for Fab domains other than trastuzumab. However, for glutamate the correlation of the excipient–residue contacts in omalizumab is lower between the two force fields (*r* = 0.46), which can again partly be attributed to the smaller value range of the glutamate contacts compare to the arginine contacts. Nevertheless, while the specificity of glutamate– residue contacts at the individual residue level is only partially captured by Martini 3-exc, the average number of glutamate contacts (averaged over all omalizumab surface residues) is 0.19, which is in reasonable agreement with the value of 0.12 from the CHARMM36m-NBF simulation (Figure S4). This result suggests that, at the global level, the description of the protein interactions with both excipients, arginine and glutamate, is improved in the Martini 3-exc model.

### General Discussion

Taken together, the new Martini 3-exc parameter set for excipient–residue interactions improves the agreement with the all-atom simulations. The arginine contacts observed in the validation system (omalizumab Fab domain) match the reference data. Substantial improvements of protein–excipient interactions are expected for Fabs of the IgG1 subclass in general. For glutamate excipients, the new Martini 3-exc parameters yield improved molecular contacts on a global average level, but contacts of a given specific residue cannot be reliably predicted by the CG force field. However, this result has to be carefully interpreted in light of the overall small number of glutamate–Fab contacts.

Many biopharmaceutical formulations include excipients that have, e.g., amide, ester, or ether groups, etc., which are chemically distinct from the amino acid excipients studied in this work. In principle, we expect the CG force field parametrization approach presented here to be generally applicable to protein interactions with excipients or ligands that have attractive interactions with the biomolecules of interest. Molecules that have less favorable (or even repulsive) interactions with the proteins, such as glutamate, are less robust because they are subject to increased statistical noise and are more sensitive to perturbations, and thus small changes of the environmental conditions or the parameters can have relatively large consequences.

One principal limitation of our approach is that inaccuracies of the all-atom force field will be transferred to the CG model. This may, for example, concern the description of the charge–charge interactions, which can be overestimated in nonpolarizable force fields with fixed point charges.^60^ Polarizable force fields may provide an improved description of such effects, but they are computationally more demanding, and they have their own limitations. A computationally tractable alternative could be the use of the electronic continuum correction (”charge-scaling”) in fixed point charge force fields.^60,61^ However, experimental data are necessary to ultimately validate the accuracy of the simulation predictions. For the systems of interest in the present study, these could for example be NMR experiments to investigate, at the level of the single amino acid residues, the protein–excipient interactions via chemical shift perturbations. However, such experiments are challenging due to the large size of the Fab domains, and such data are currently not available.

A popular strategy to improve protein–protein interactions in Martini, first suggested by Stark et al. for Martini 2, is to globally scale the LJ well-depths of all protein–protein bead interactions.^62^ In Martini 3, Thomasen et al. obtained improved agreement with experimental SAXS and NMR data either by globally downscaling the protein–protein interactions (less attractive) or by upscaling the protein-water interactions (more attractive).^41,42^ In the present study, for protein–excipient interactions, we found that such global scaling is not sufficient. Instead, we observed that the favorable interactions of arginine and glutamate with Arg, Asp, Glu, Lys, Phe, and Tyr residues need to be increased, while decreasing the interaction strength with all other residue types (except for glutamate excipients with Ile) was necessary to obtain improved results. For full-length antibodies, which also include the Fc domain, the parameters derived here should provide more accurate protein–excipient interactions, but additional refinement may be necessary. This is challenging also in light of the glycosylation of the Fc domain, and would also require to further refine Martini 3 carbohydrate parameters^63–65^ for glycosylated amino acid side chains. However, once these are available, the parametrization strategy followed in this work can be promising also for the parametrization of excipient–glycan interactions.

## Summary and Conclusions

The Martini force field is a computationally efficient coarse-grained model that is promising for studying biomolecular systems on time and length scales that are currently out of reach for all-atom simulations. By construction, the parameters of CG force fields can be expected to be transferable to only within a narrow range (and their applicability within that range needs to be carefully tested), and thus systems or conditions that differ substantially from those used in the original parametrization are usually not as accurately described with the default parameters. However, the modular structure of the Martini force field facilitates adaptations and optimizations for new systems and compounds, as this work demonstrated for protein–excipient interactions as a case example.

In the present work, we introduce an adapted Martini 3-exc coarse-grained force field of the amino acid excipients arginine and glutamate, which was parametrized against all-atom simulations as a reference. To study the effects of excipients on aggregation and solubility of antibody Fab domains at a near-atomic level, accurate excipient–residue interactions are crucial. For this reason, the Fab domain of the monoclonal antibody trastuzumab was used in excipient–Fab simulations to compare the local excipient–residue interactions, using the new CG Martini 3-exc excipient models described in this work. The excipient–residue interaction parameters were reparametrized to match molecular contacts of reference simulations using the CHARMM36m-NBF all-atom force field. The new Martini 3-exc was validated with the omalizumab Fab domain, which was not used in the parametrization procedure.

The adjusted excipient–residue interaction parameters of Martini 3-exc yield significantly improved results for the interactions of the trastuzumab Fab domain with arginine and glutamate excipients. The validation simulations with the omalizumab Fab domain (which was not used in the parametrization procedure) showed that for arginine excipients the new interaction parameters of Martini 3-exc provide results in agreement with the all-atom reference simulations, but for glutamate the Fab–excipient contacts were only slightly improved compared to the original Martini 3 force field.

We expect the adopted parametrization strategy to be applicable also to other excipients or ligands that have attractive interactions with proteins. For proteins that are very different from the ones used in this study, the refined Martini 3-exc parameters should be carefully checked and validated, and further refinements may be necessary.

In conclusion, the Martini 3-exc parameter set provides improved descriptions of excipient– Fab interactions, opening the way towards studying the molecular underpinnings of the stabilizing effects of arginine and glutamate excipients on the solution state properties of biopharmaceutical high-concentration monoclonal antibody formulations using CG-Martini simulations.

## Data and Software Availability

All relevant data are included in the Tables and Figures provided in the manuscript and Supporting Information. MD simulations were performed with GROMACS version 2021.1 (https://manual.gromacs.org/documentation/2021.1/download.html). The new Martini 3-exc force field parameter files, starting coordinates of the MD simulations, MD parameters file, and force field topology files are enclosed in GROMACS format as Supporting Information.

## Supporting Information Available

Correlation of molecular contact cutoff of 0.50 nm and 0.75 nm in coarse-grained and all-atom resolution. Histograms of bonds and angles of the 4-bead arginine model after reparametrization compared to the all-atom histograms. Histograms of bonds and angle of the 3-bead glutamate model after reparametrization compared to the all-atom histograms. Arginine contacts with trastuzumab residues in Martini 3, Martini 3-exc and CHARMM36m-NBF. Correlation of molecular contacts between arginine and glutamate with omalizumab in Martini 3 compared to CHARMM36m-NBF.

Martini 3-exc force field parameter files (ZIP) as well as coordinate files with starting coordinates of the simulations, MD parameters (MDP) file, and force field topology files (ZIP).

## Acknowledgement

The authors thank Felicitas Schmadl and Marina Rechsteiner for technical support and Raphael Drerup for project administration support. This work was supported by Deutsche Forschungsgemeinschaft (DFG) under Germany’s Excellence Strategy - EXC 2033 – 390677874 - RESOLV. This project was carried out in preparation of SCHA 1574/8-1.

## Notes

### Competing Interest Statement

The authors have declared no competing interest.

### Summary of Updates

Method section updated with more computational details; Typo in Figure 2 fixed; Results and Discussion section updated with discussion about accuracy of all-atom force fields.

