## Supplemental Figures for "Optimized Protein–Excipient Interactions in the Martini 3 Force Field"

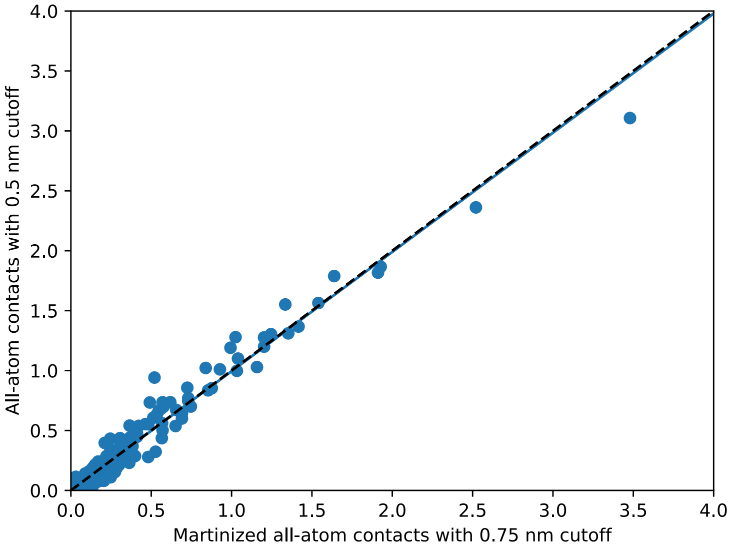


Figure S1: Correlation plot of arginine contacts to trastuzumab Fab residues with 0.5 nm cutoff in all-atom simulation versus contacts in the same simulation analyzed in terms of its CG-Martini 3 coordinates with 0.75 nm cutoff. The dashed black line is the diagonal ($y = x$). The Pearson correlation coefficient is 0.98 and the slope of the linear fit (blue line) is 0.995.

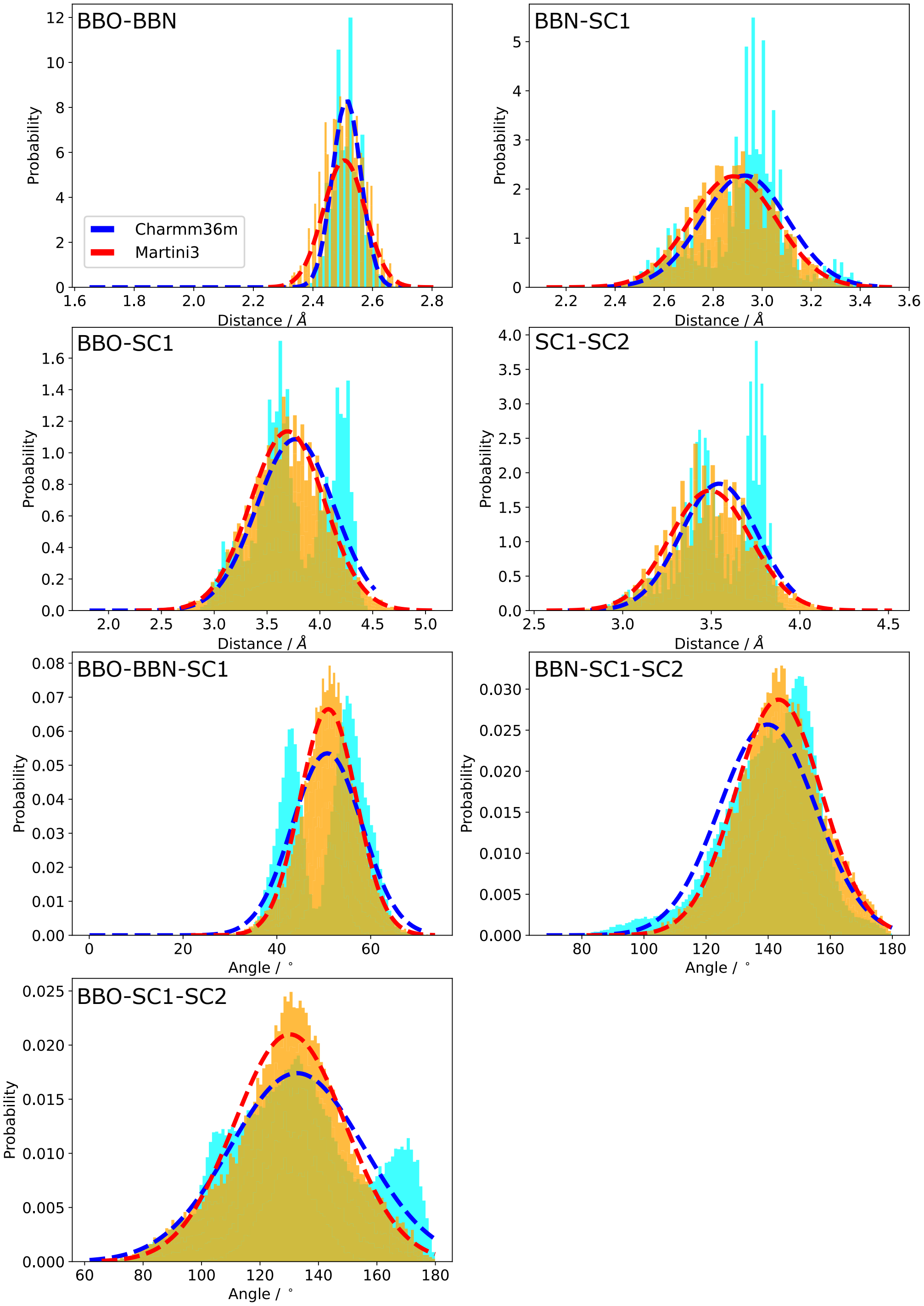


Figure S2: Histograms of intramolecular distances and angles of the Martini 3 arginine excipient molecules compared to the all-atom reference. BBO and BBN are the SQ5n and SQ4p CG beads for the backbone carboxyl- and amino-groups, respectively, and SC1 and SC2 refer to the side chain beads. The bimodality of some of the all-atom distributions cannot be captured by the harmonic potentials used in the CG force field, which give rise to Gaussian distributions.

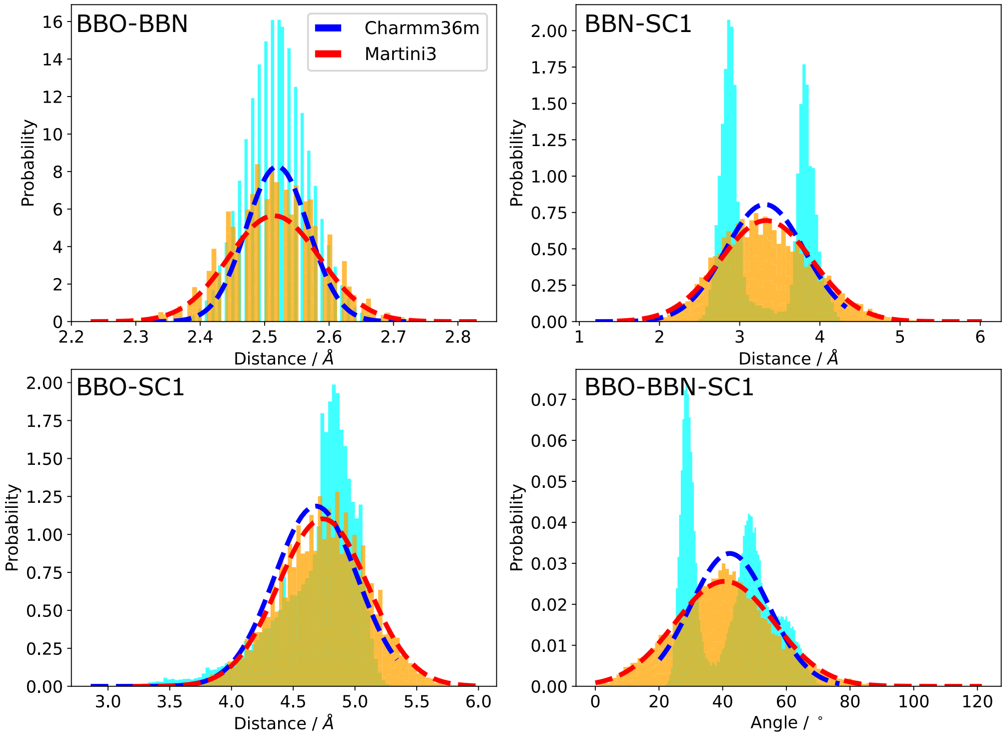


Figure S3: Histograms of intramolecular distances and angles of the Martini 3 glutamate excipient molecules compared to the all-atom reference. BBO and BBN are the SQ5n and SQ4p CG beads for the backbone carboxyl- and amino-groups, respectively, and SC1 refers to the side chain bead.

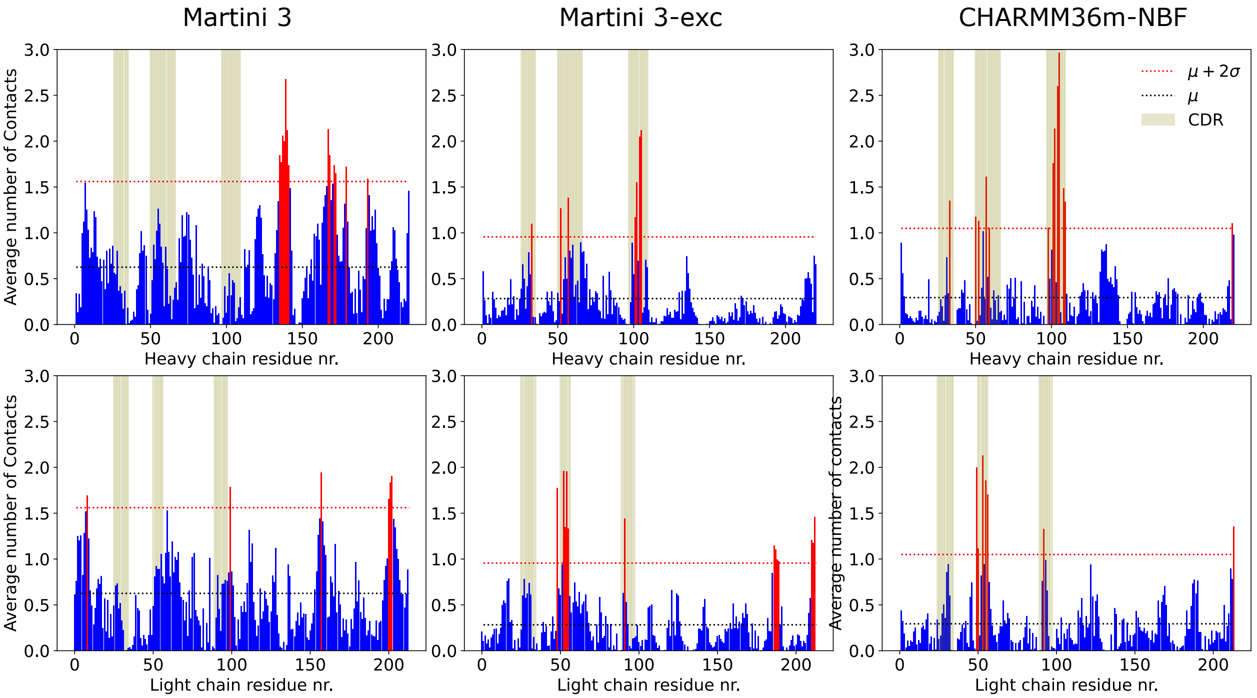


Figure S4: Comparison of arginine–trastuzumab contacts in the Martini 3, Martini 3-exc, and CHARMM36m-NBF simulations. The CDR regions are highlighted by light green bars. The contact count averaged over all residues is shown as a black dotted horizontal line. The residues marked in red indicate the amino acids whose arginine contacts are greater than the average by more than two standard deviations (dashed red horizontal line).

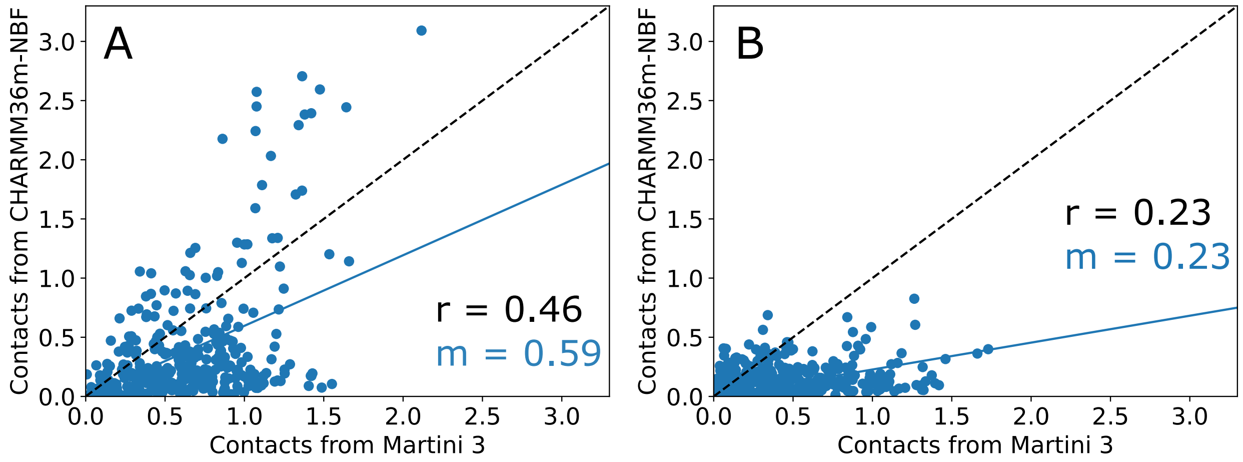


Figure S5: Correlation of protein–excipient contacts between CHARMM36m-NBF and Martini 3 simulations of the omalizumab Fab domain (r denotes the Pearson correlation coefficient between the two data sets, m is the slope of the linear fit (blue line)). The arginine-residue and glutamate-residue contacts are depicted in panel A) and B), respectively.
